# 3D-printed toolkit for regulating epithelial unjamming

**DOI:** 10.1101/2025.11.13.688246

**Authors:** Yael Lavi, Yehuda Sandler, Areej Saleem, Liav Daraf, Shahar Nachum, Lior Atia

**Affiliations:** Department of Mechanical Engineering, Ben-Gurion University of the Negev, Beer-Sheva 8410501, Israel

## Abstract

During wound healing and cancer invasion, epithelial monolayers transition from a solid-like, jammed state to a fluid-like unjammed state. However, affordable and accessible tools for modeling and precise tuning of this response remain limited. Here, we present a simple, lithography-free 3D-printed toolkit of customizable inserts and scratchers for fabricating micro-gaps with defined geometries and widths down to ∼50 µm, without requiring cleanroom facilities or specialized expertise. Using epithelial monolayers, we demonstrate that gap width fundamentally regulates unjamming dynamics. Narrow gaps induced rapid, coordinated migration with high initial velocities, whereas wide gaps suppressed early unjamming and produced slower, spatially heterogeneous closure. Cell shape dynamics confirmed conserved elongation, cell rounding, and closure behaviors across conditions. The presented toolkit provides new opportunities for investigating the mechanical and molecular mechanisms that regulate epithelial plasticity, enabling fine control of unjamming transitions and allowing the modeling of both healthy and pathological wound-response regimes.

## Introduction

The surfaces of organs and cavities are lined by cohesive epithelial sheets. These tissues are composed of tightly packed cells that resist motion and collectively behave as a solid-like jammed matter. Yet during key physiological events in embryogenesis, wound healing, and cancer invasion, epithelial cells can adopt a fluid-like state and unjam.^1-21^ This transition, known as cellular unjamming, has emerged as a central theme in the field of tissue mechanics.^12-21^

The framework of collective epithelial jamming was adopted from the field of condensed particle systems, like compacted grains and sand.^14,19,22^ In inert particulate system, promoting unjamming is relatively simple and can be achieved by reducing particle density.^22,23^ In epithelial systems, promoting unjamming is not trivial, as cells actively remodel their cytoskeleton and adjust their adhesion proteins.^4,7-9,15^ Thus, simple and accessible tools that can promote epithelial unjamming in a controlled manner are needed to enable deeper investigation of the underlying mechanisms of these transitions. Our aim is to provide a simple and affordable tool for promoting epithelial unjamming by generating a controlled and narrow gap within a confluent monolayer. A gap that is just a few cell widths wide can provoke local migration, transient unjamming, and eventual re-jamming as the gap closes, as illustrated in Figure 1A. Existing methods for producing controlled, narrow gaps rely on complex and costly microfabrication methods and require specific technical expertise. These include laser ablation,^24,25^ ion etching,^26^ digesting microflows,^27^ robot-assisted wounding,^28^ and photolithography-based techniques.^29,30^ Such methods are often inaccessible to most molecular biology labs.

**Figure 1.**
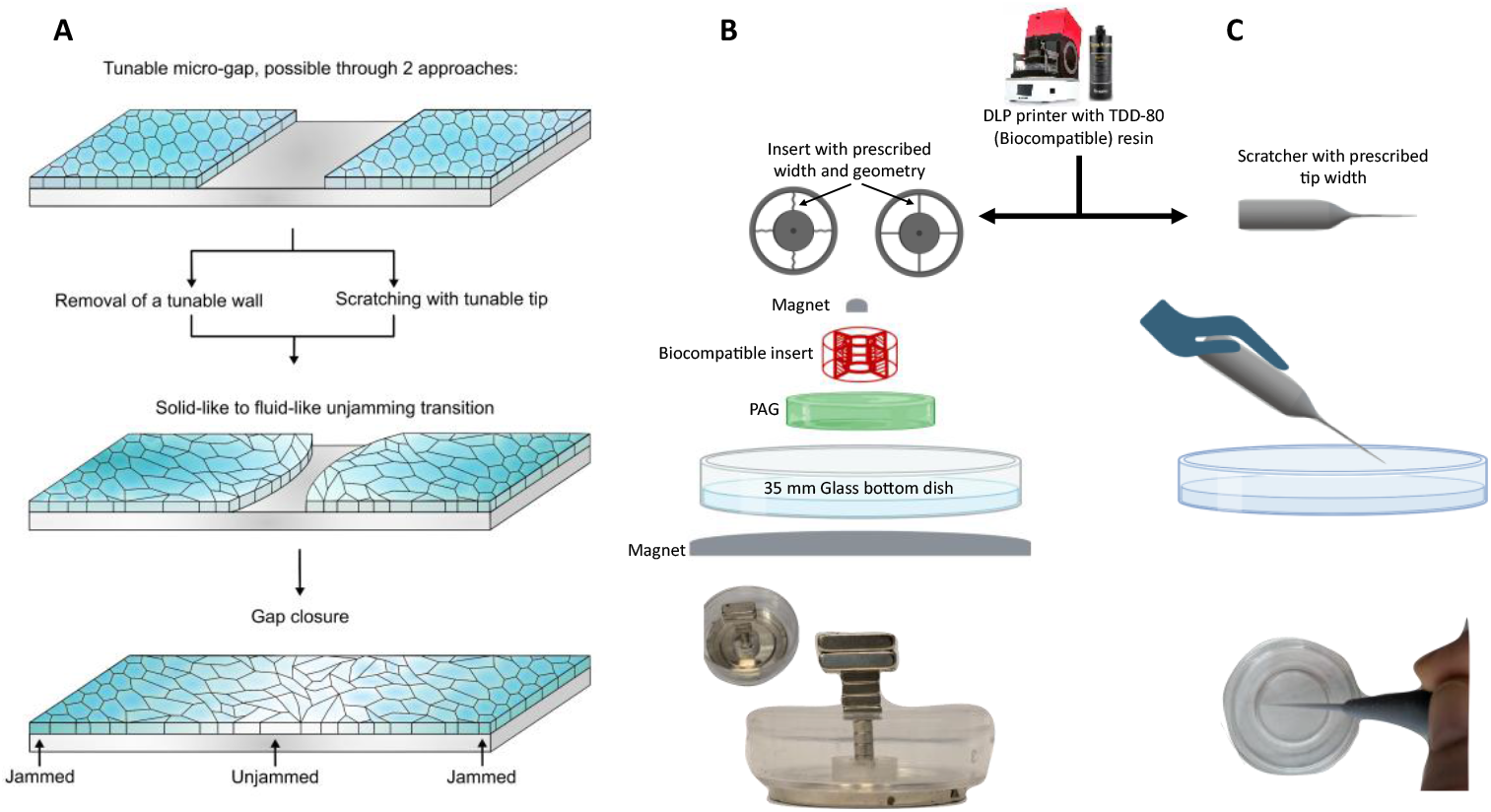
Schematic overview of a 3D-printed method for generating tunable epithelial micro-gaps using inserts and scratchers, enabling controlled unjamming. (**A**) An illustration of a controlled gap within a confluent monolayer, followed by the onset of collective migration as cells transition from a solid-like jammed state to a fluid-like unjammed state during gap closure. (**B**) Integration of biocompatible, 3D-printed inserts with a Polyacrylamide gel (PAG) substrate, and pressing magnets for sustained seal. The entire setup fits into a 35 mm glass-bottom dish, enabling easy fabrication, high reproducibility, and improved sealing for creating micro-gaps between confluent epithelial monolayers. (**C**) A custom-designed, adjustable-width scratching tool allows controlled removal of cells, generating micro-gaps of defined widths. The correlation between the degree of unjamming and gap size, as revealed here, demonstrates the ability to control the extent of unjamming.

We present a simple, compact, lithography-free Digital Light Processing (DLP) approach for generating tunable, clear, and cell-free micro-gaps, using resin-based 3D printing. This technology, which is increasingly used in biomedical research^31-33^ and the dental industry,^34,35^ enables rapid fabrication of smooth, high-resolution structures without the need for clean rooms, lasers, special coatings, or specialized expertise. Similar to other 3D-printing technologies^36,37^ it is also widespread in shared academic facilities and by online vendors.

Here, we developed a toolkit to regulate epithelial unjamming using a DLP printer with a biocompatible resin. The printer projects an image onto a vat of liquid photopolymer, solidifying it layer by layer to form precise 3D objects (Figure 1B, C). We produce customizable 3D-printed micro-scratchers and inserts that can generate reproducible micro-gaps in epithelial monolayers. Our approach enables rapid fabrication, enabling fine control of gap morphology, which is rarely achievable in conventional scratching or barrier-based assays.^24-30,38-41^ Furthermore, this method enables the generation of co-culture systems while substantially reducing cell mixing, a limitation commonly encountered in double-seeding or backfilling approaches.^42^ The toolkit enables precise control over the unjamming response, as evidenced by its correlation with the micro-gap size. It provides a robust, highly versatile, simple, and affordable (approximately 3 cents per insert or shaped islet and 30 cents per scratcher) tool for investigating the underlying mechanisms of unjamming transitions.

## Results

### A toolkit for controlled unjamming

We designed two separate tools for generating micro-gaps in epithelial monolayers: (1) removable inserts and shaped islets with variable widths and edge geometries (Figure 1B), and (2) reusable scratchers with different tip widths (Figure 1C). Both the inserts and the scratcher were designed in SolidWorks and fabricated on a 3D DLP printer with a biocompatible resin. This method does not require any post-processing or surface treatment prior to cell seeding, thereby further streamlining the preparation process. To demonstrate the abilities of this toolkit, we used Madin-Darby Canine Kidney, type II (MDCK II) cells throughout the study.

Inserts were fabricated with varying widths and geometries (Figure 2, rows 1-2, columns 1-2), enabling both linear and non-linear gap formation. The latter are particularly valuable, as *in vivo* geometries are often curved, branched, or irregular rather than strictly linear.^43-45^ Beyond generating gaps, our system can fabricate cell islets of controlled shape. These islets are discrete regions surrounded by printed walls, enabling precise spatial patterning of cell populations (Figure 2, row 3). Cells reliably adhered, spread, and proliferated within these predefined geometries, recapitulating fine structural features down to ∼ 50 µm scale (Figure 2, column 4). Furthermore, the system is readily adaptable for co-culture applications, enabling spatially controlled seeding of distinct cell populations without intermixing.^42^ This is demonstrated with cells stably transfected to express either ZO1-EGFP^46^ or the FUCCI transgene^47^ (Figure S1). Lastly, our toolkit also provides customized scratchers to inflict a micro-wound upon a confluent layer, leaving a clear and cell-free micro-gap (Figure 2, row 4).

**Figure 2.**
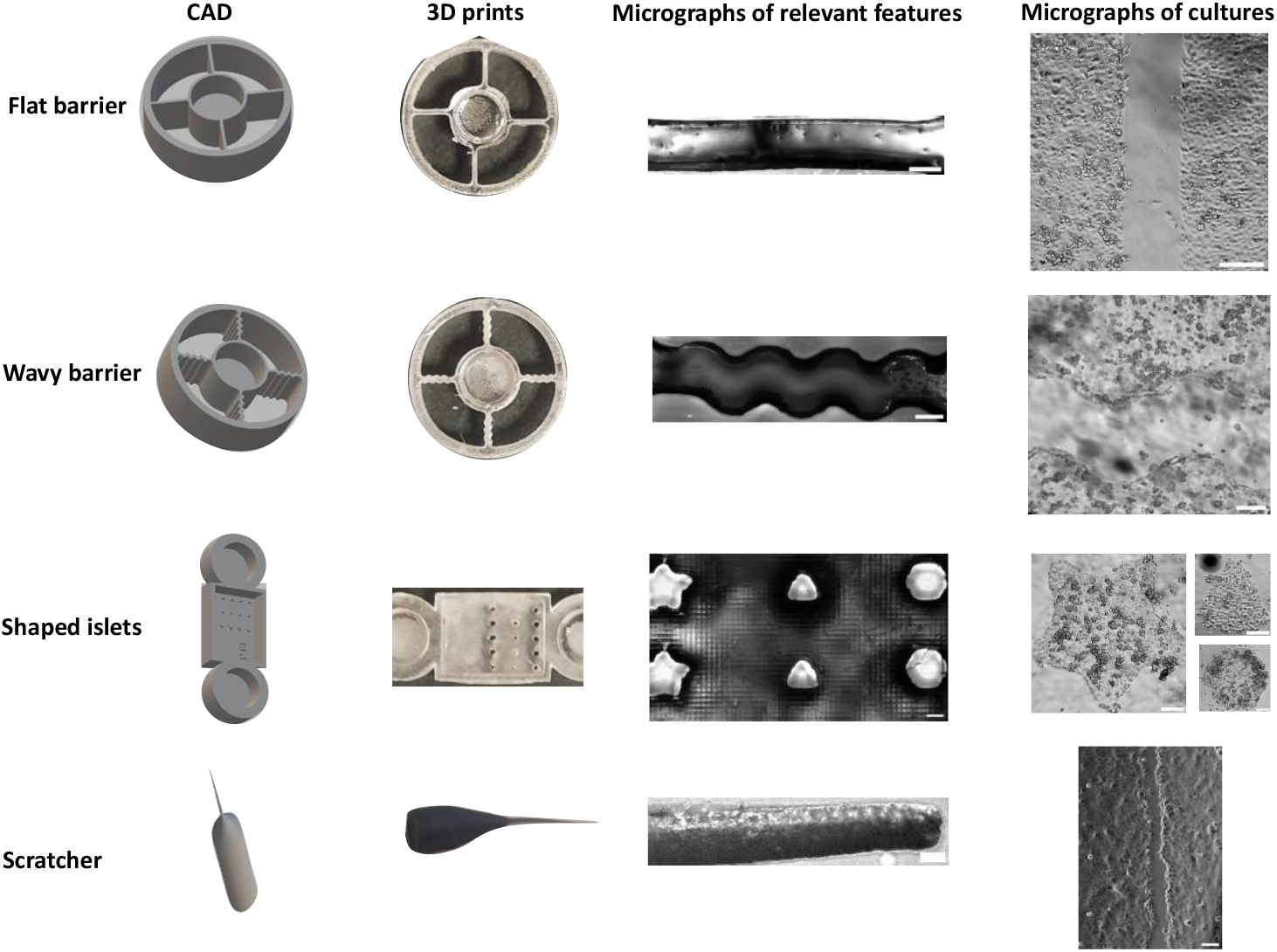
Versatility of the 3D-printed toolkit for generating controlled epithelial microenvironments. Each row represents the progression from a CAD model (left; supplementary files) to printed parts, structural micrograph, and the resulting cell culture (right). **Top row**: Flat barrier design produces linear micro-gaps of defined width, supporting uniform epithelial monolayers and clean exclusion zones. **Second row**: Wavy barriers introduce curvature into the gap, enabling the study of geometrical effects on unjamming and closure dynamics. **Third row**: Shaped islets of various geometries (triangular, star-like, hexagonal) enable shape-dependent behavioral assays of epithelial monolayers. **Bottom row**: Custom-designed scratchers with tunable tip widths enable generating narrow micro-gaps by manual scratching. Scale bars: 100 μm.

### Insert-based assays

To quantitatively evaluate how the initial gap width influences collective migration dynamics, we seeded cells in inserts with wall widths of 50, 100, or 250 µm. A key technical challenge was preventing cells from growing underneath the insert walls. To address this, we developed a reliable magnet-assisted sealing strategy: polyacrylamide gel (PAG) was cast onto the glass bottom of a petri dish and subsequently coated with collagen to promote cell adhesion. The insert was then placed directly on the treated PAG. To ensure tight contact between the insert and the gel, magnets were positioned above the insert and beneath the culture dish, generating a compressive force that sealed the insert against the gel (Figure 1B, magnets model). This configuration effectively excluded cells from underneath the wall region, producing consistent and reproducible gap geometries (Figure 2, rows 1-3, column 4).

Unlike PDMS-based or molded stencil systems^38^, this approach requires no adhesives, surface coatings, or alignment steps. Once confluence was reached within flat-wall inserts, they were removed, allowing cells to migrate into the newly exposed, cell-free region. We compared the migratory behavior across the three gap widths (Figure 3) by quantifying velocity and cell shape metrics from time-lapse images that followed the two advancing layers to their convergence.

**Figure 3.**
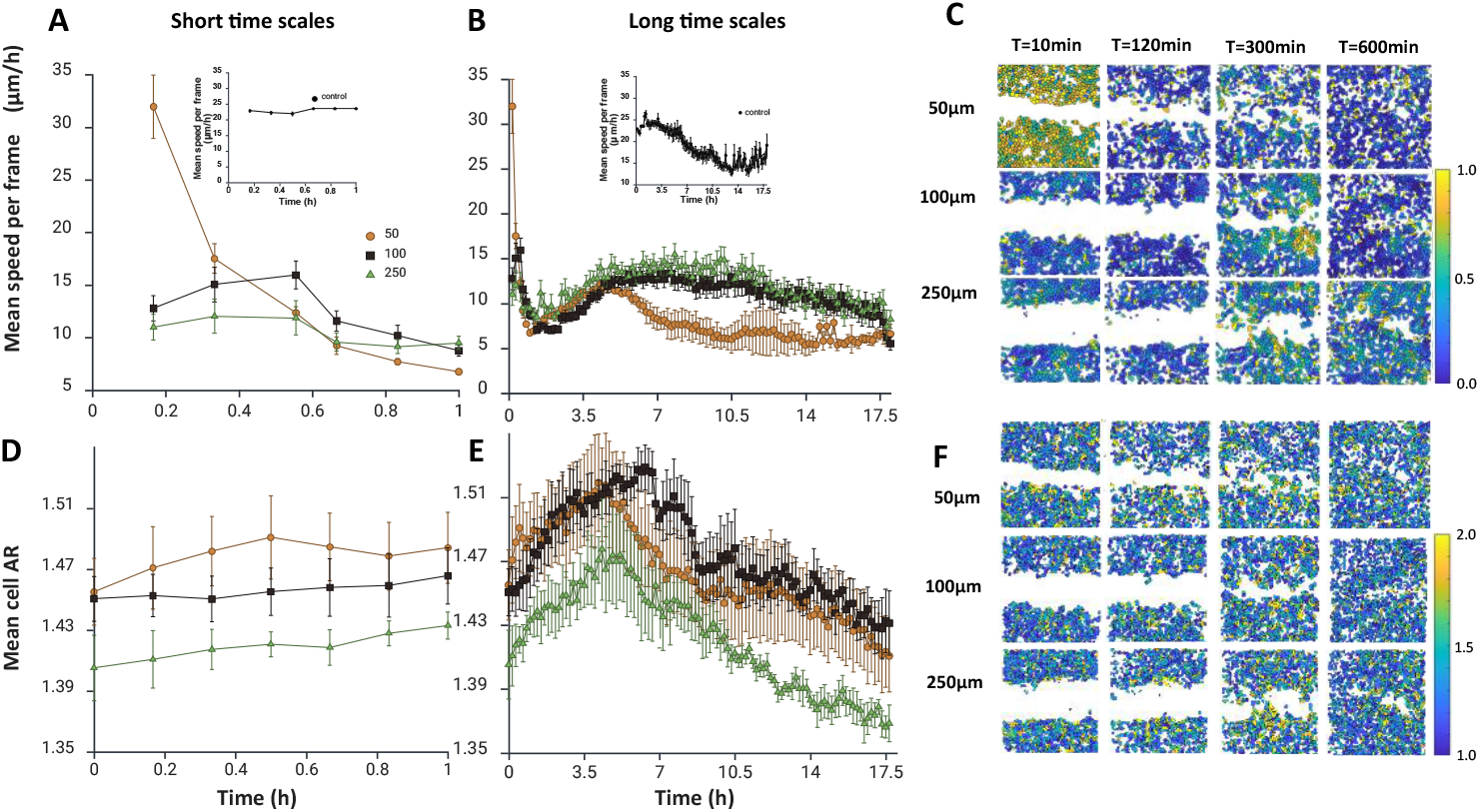
Insert-based assays reveal that wide gaps tend to inhibit cellular motion, while narrow gaps promote it. (**A, B**) Time course of mean velocity for epithelial sheets, closing gap widths of 50 µm (•), 100 µm (◼), and 250 µm (▲) (mean ± SEM, n = 4, 7, 5, respectively). On short time scales (**A**), narrow gaps trigger a velocity peak higher than the control (inset; no removal of insert), while wider gaps initially inhibit cellular dynamics, indicated by significantly lower velocities. On longer time scales (**B**), a second velocity peak was followed by a rapid decline in narrow gaps, whereas the decline in wider gaps was significantly delayed. (**C**) Spatial heatmaps of velocity magnitude at 10, 120, 300, and 600 min after barrier removal. Rows correspond to initial gap widths. At *t* =10 min, narrow gaps show fast coordinated motion, while wider gaps show isolated pockets of slower velocity. Color bar indicating normalized mean velocities. (**D, E**) Time course of mean cell aspect ratio (AR) for epithelial sheets closing the gap (mean ± SEM, n = 4, 7, 5, respectively). Elongation, marked by an increasing AR and indicating unjamming, peaks concurrently with velocity, followed by cell rounding, marking the return to a jammed state. (**F**) Corresponding spatial maps of AR at 10, 120, 300, and 600 min after barrier removal. Color bar indicating AR. Note that white patches within the monolayers are a result of miss identification.

On short time scales (up to 1 h), wide gaps (100 and 250 µm) substantially inhibited migratory dynamics compared with the control (Figure 3A and inset). In contrast, narrow gaps (50 µm) elicited significantly higher initial velocities. We note that insert removal and microscope setup required ∼ 15 min; therefore, the earliest migration phase could not be captured. Velocities declined rapidly thereafter across the three widths. On longer time scales, the early velocity drop was succeeded by a slower rise, peaking at around 5 hours for all three widths (Figure 3B). Following this rise, the velocity steadily decreased for narrower gaps, whereas wider gaps exhibited a prolonged plateau phase before a subsequent decrease in velocity. For narrower gaps, a complete closure coincided with the velocity peak at 5 hours. The rapid decline in velocity thereafter represents the re-jamming of the layer post gap closure (Movie S1). For wider gaps, the velocity peak at 5 hours marked the onset of closure, as these gaps promoted slower, spatially staggered dynamics, reminiscent of a zipper-like fastening closure (Movie S2). Once closure was completed, velocities decreased as the monolayer re-jammed. This dynamical heterogeneity was clearly visible in the velocity heatmaps (Fig. 3C).

With respect to cell dynamics narrower gaps yielded faster closure and higher peak velocities, whereas wider gaps induced a delayed and heterogeneous migratory response. With respect to cell geometry, mean aspect ratio (AR), a proxy for cell elongation, increased early during migration (Figure 3D), peaking concurrently with the velocity maximum in both the 50 µm and 250 µm conditions (Figure 3E). For narrow gaps, AR heatmaps show that cell elongation occurs at all distances from the edge, while for wider gaps it tends to occur closer to the edge (Figure 3F). Increased AR is a hallmark of unjamming transitions, reflecting cytoskeletal polarization and loss of positional confinement.^12-21^ Interestingly, the rates of AR increase and decrease were similar across all conditions, indicating conserved elongation and rounding processes regardless of gap width.

The significantly lower starting velocity and AR in the 250 µm condition indicate that the monolayer at *t* = 0 was in a more tightly jammed state compared with narrower gaps. In parallel, mean cell area decreased over time for all gap widths (Figure S2A). This suggests that unjamming occurred regardless of layer maturation and continuous proliferation when using inserts.

The results from the insert experiments imply that the presence of a wall near the cells, prior to insert removal, inhibited the initial cellular dynamic response immediately after insert removal. The exception was the narrowest gap (50 µm), which prompted us to consider whether the close proximity between the layers evoked a gradient-dependent cellular mechanism, such as collective chemotaxis,^48^ that could overcome this inhibition. Although chemotaxis was not examined in this study, its potential involvement represents an interesting direction for future investigation.

### Scratch-based assays

To evaluate the scratch-based assay, we used our custom scratcher to generate wall-free micro-wounds in confluent monolayers. The resulting gaps were clean and reproducible, though minor width variations arose from differences in applied manual force during scratching. Gaps of varying widths were analyzed as in the insert-based assays analysis.

At short time scales, all gaps elicit an initial velocity that is higher (Figure 4A) than control (Figure 3A, inset). The narrower the gap, the greater the initial velocity. Following this initial rise, velocities quickly decline across all conditions, resembling the dynamics observed in the 50 µm insert-based gaps and contrasting with the behavior of the wider (100 and 250 µm) insert-based gaps. These results further support the notion that wide insert-based gaps restrain unjamming dynamics.

**Figure 4.**
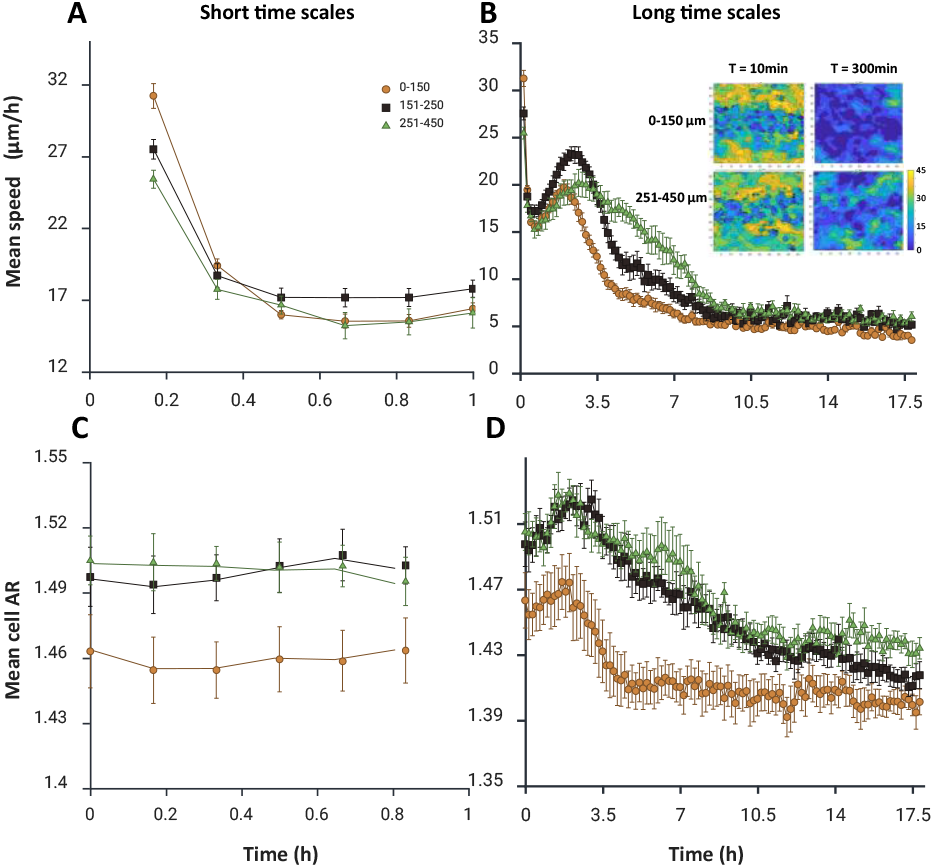
Scratch-based assays triggered coordinated motion whose extent scaled with gap width. **(A, B)** Mean velocity of epithelial monolayers migrating to close gaps of varying initial widths: 0–150 µm (•), 151–250 µm (◼), and 251–450 µm (▴) (mean ± SEM; n = 7, 5, 4, respectively). On short time scales (**A**), velocity starts higher than the control (Figure 3A, inset) for all scratch widths, with narrower scratches exhibiting higher initial velocity. On longer time scales (**B**), velocity first declines before peaking. Narrow scratches show velocities that peak at 3.5 h, while wider scratches peak progressively later. The inset shows spatial velocity heatmaps, at *t* = 10 min and 300 min, and for 0-150 µm and 251-450 µm wide scratches. Color bar indicates mean velocity by PIV analysis. These maps show that cellular velocity was coordinated at *t* = 10 min for both widths (see also supp. movies S3 and S4). By t = 300 min, narrow scratches showed markedly diminished and plateaued dynamics, whereas wider scratches retained higher velocities. Velocities subsequently declined for all widths, with narrower scratches exhibiting faster declines. (**C, D)** Mean AR of monolayers migrating to close a gap of 0–150 µm (•), 151–250 µm (◼), and 251–450 µm (▴) (mean ± SEM; n = 7, 5, 4, respectively). AR increased marking elongation and unjamming. On longer time scales (**D**), elongation peaked concurrently with velocity peaks, then declined rapidly for narrow scratches, and more gradually for wider ones.

At long time scales, velocities rose, peaking earlier than in insert-based experiments; sooner for narrow scratches and progressively later for wider ones. This progression in peak times correlated with the distance cells had to traverse. As in insert-based assays, velocities peaked in sync with gap closure. Velocities rapidly decreased across all conditions afterwards. Velocity heatmaps (Figure 4B, inset) confirmed uniform collective migration across the entire field for all scratch widths (Movies S3 and S4). Notably, overall velocities were higher for scratch-based assays than for insert-removal experiments of comparable width.

With respect to cell geometry, mean AR increased sharply after the first hour (Figure 4C-D). The inclines of all 3 graphs were similar, again suggesting a conserved elongation process. AR peaked concurrently with the velocity peak for all scratch widths. These velocity and AR peaks were in sync with the closure of the micro-gaps. Following this peak, the narrower scratches showed a sharp decline in AR, followed by a plateau at an AR of ∼1.39. Wider scratches also showed a decline, although it was less pronounced. The observed AR values are consistent with those predicted by the biophysical vertex-model simulations^1-21^ for a jammed monolayer. Cell area increased initially, marking the spreading of leading cell rows, and then declined with the convergence of the two layers (Figure S2B).

## Discussion

Here, we present a simple toolkit designed to lower the technical barrier for conducting controlled unjamming experiments. The comparative analysis of insert-based and scratch-based assays highlights the flexibility and range of migration behaviors achievable with this system. Insert-based experiments with narrow walls induced rapid, coordinated motion and higher peak velocities, while wider walls produced slower and more spatially heterogeneous migration patterns.

Scratch-based experiments, regardless of gap widths, consistently generated higher velocities and more coordinated collective motion than their insert-based counterparts. Hence, thin-wall inserts and all scratch-based assays induced qualitatively similar dynamics in trends and may serve as useful models for physiological wound responses under healthy conditions. However, thick walls inhibit the initial unjamming response, suggesting potential relevance for modeling impaired or pathological conditions. All in all, both insert and scratch-based assays showed a strong correlation between gap width and the extent of unjamming, emphasizing the tunability of the system.

Regardless of the type of assay, the initial distance between opposing cell fronts at the time of gap infliction emerged as a fundamental parameter governing the magnitude of the unjamming response. Whether this dependence arises due to collective chemotaxis^48^, or another type of mechanism, remains an open question for future endeavors. Researchers can tailor the system to specific experimental aims, using scratchers or narrow inserts for fast closure and wide inserts for slower, spatially heterogeneous migration dynamics.

## Materials and Methods

### 3D Printing of Inserts and Scratchers

We utilized the Asiga MAX 3D printer to fabricate the inserts and scratchers. Designs were created in SolidWorks and exported as STL files. These files were then imported into Asiga Composer, the printer’s build-preparation software, where parts were arranged and duplicated as needed. To ensure optimal surface quality, we oriented the parts so that the surfaces requiring smoothness faced the printer’s screen, away from the polymerization plate. For printing, we selected the TDD-80 clear resin, known for its clarity and biocompatibility. The resin was poured into a dedicated vat and stirred thoroughly to eliminate any pre-polymerized remnants from previous prints. Air bubbles introduced during stirring did not affect the printing process. The TDD-80 resin requires preheating to 30 °C, which we allowed to proceed for optimal results.

When printing was completed, the polymerization plate rose with the printed parts adhered to it. We then allowed a few minutes for excess resin to drain back into the vat, assisting the process by gently running a spatula around the plate. Once drainage was sufficient, we removed the plate and placed it into a shaking bath filled with Isopropyl Alcohol (IPA) for 3 minutes. The wash was repeated with fresh IPA. After washing, we detached the parts from the polymerization plate and placed them into a beaker of IPA, manually agitating for 1–2 minutes. This was especially crucial for parts with printed open cavities that could retain trapped and unpolymerized resin. Following agitation, the parts were left to dry completely. Once dry, we gently cleaned any excess resin using a small, slanted tweezer wrapped in absorbent paper. This step was vital to prevent deformation during curing and to avoid introducing potential toxicity into cell cultures. The cleaned parts were then cured under UV light for 1 hour on each side. If the inserts became too brittle, we reduced the curing time accordingly.

### Preparation of Polyacrylamide Gel (PAG) Islets

To facilitate sealing, we used magnetic forces to gently press down a printed insert against a Polyacrylamide Gel (PAG) substrate (Figure 2B). Working with PAG substrates also allowed control over substrate rigidity and ECM coating, while providing the future option to incorporate traction force microscopy for mechanical force measurements and still produce microgaps.^40^ The gels were formed by depositing the gel solution onto the glass bottom of a Petri dish and covering it with a coverslip, ensuring a smooth surface. To promote adhesion of the PAG to the dish, we first activated the dish surface. Activation was performed under a chemical hood. An activation solution was prepared by mixing 8.572 ml of 99% ethanol (Romical), 714 µl of acetic acid (Bio-Lab), and 714 µl of 3-(Trimethoxysilyl)propyl methacrylate (bind silane, Sigma). Each 35 mm glass-bottom Petri dish was deposited with 900 µl of the solution and left for 10–30 minutes, at room temperature, under a chemical hood. Dishes were then washed twice with 99% ethanol and dried using compressed air.^49^

To prevent premature polymerization, we prepared the gel solution only after dish activation. Alternatively, we prepared the solution without initiators and kept it on ice, adding initiators immediately before application. For ∼ 500 µl of PAG solution, we combined 310 µl PBS (Diagnovum), 150 µl of 40% acrylamide solution (BIO-RAD), and 37.5 µl of 2% bis-acrylamide solution (BIO-RAD). Just before application, we added 2.5 µl of Ammonium Persulfate (APS, BIO-RAD) and 0.25 µl of TEMED (Sigma), mixing gently to avoid air bubbles.^49^ To create PAG substrates, we deposited 20 µl of the gel solution near the edge of each activated dish’s inner well. A 16 mm diameter coverslip was placed over the gel droplet, allowing it to spread evenly. Gels were left to polymerize for 1 hour at room temperature, under a chemical hood. Post-polymerization, 1 ml of PBS was added to each dish, and coverslips were gently removed using a scalpel or fine forceps. For curing and sterilization, dishes containing polymerized PAG substrates in PBS were placed under UV light in a biological hood for 20 minutes. Subsequent steps were conducted under sterile conditions in the biological hood.

### PAG Surface Activation and Coating

To promote cell adhesion, we activated the PAG surface using the crosslinker sulfo-SANPAH (Moshe Stauber Biotec Applications), which facilitates the binding of extracellular matrix proteins like collagen (PuerCol, Advanced BioMatrix) to the gel surface. Sulfo-SANPAH was reconstituted by adding 1 ml of PBS to 100 mg of the SANPAH powder, preparing aliquots of 20 µl in dark Eppendorf tubes, and storing them at −80 °C. Prior to use, aliquots were thawed and diluted with 480 µl PBS to obtain 500 µl of working solution. Each PAG-containing dish received 70–100 µl of this solution. Dishes were then exposed to UV light for 7 minutes to activate the crosslinker. We note that SANPAH should be thawed and diluted just before use, since it’s only active for a short while. While SANPAH was activating, we prepared a collagen solution at a 0.1 mg/ml concentration in HEPES (Diagnovum) buffer. For instance, to prepare 5.5 ml of this solution, we mixed 183 µl of a 3 mg/ml collagen stock with 5.317 ml of HEPES buffer. After UV activation of the Sulfo-SANPAH, dishes were washed thoroughly with PBS until the orange hue of SANPAH was no longer visible. Subsequently, 1 ml of the collagen solution was added to each dish, and the dishes were incubated at 4 °C overnight.

### Assembly and Cell Seeding

The following day, collagen-coated dishes were washed twice with PBS and left open in a biological hood to dry slightly. Inserts were sterilized by immersion in 70% ethanol for 5 minutes and then dried on absorbent paper or with compressed air. To ensure a secure seal between the insert and the PAG, we employed a magnetic system. Each setup included a large magnet placed beneath the dish and several small magnets positioned in designated wells within the inserts (Figure 1B).

We cultured MDCK II cells in EMEM supplemented with 10% FBS, 1% L-glutamine, and 1% penicillin-streptomycin, maintaining them at 37 °C with 85% humidity and 5% CO_2_. Cells were trypsinized for 10 minutes and centrifuged at 800 rpm for 5 minutes. During cell preparation, we assembled the dish by placing the large magnet beneath it, aligning the insert over the PAG, and securing it with the small magnets (see Figure 1 for full setup). Cells were resuspended at a concentration of 1.5 million cells/ml, and the appropriate volume was seeded into each well of the insert. Dishes were incubated for 1 hour, after which the medium was replaced, followed by an additional 4-hour incubation. Just before imaging, inserts were carefully removed to prevent premature cell migration.

### Imaging and Image Analysis

Imaging was performed with an Axio Observer 7 (ZEISS) equipped with a 10x Plan-Apochromat objective (ZEISS). Phase-contrast images were captured with 10-ms exposure, and fluorescence images (EGFP/mOrange) were acquired with matching filter sets (50/43, ZEISS) with 150-ms exposure. Phase images were segmented with Cellpose after training a custom model.^50,51^ The segmented images were then analyzed using TrackMate and TPIV plugins in FIJI.^52,53^ To extract and visualize analysis results, we used in-house Python and MATLAB codes. Figures were created in BioRender.

## Acknowledgments

We thank and acknowledge the helpful discussions with all members of the Atia lab, as well as the preliminary research conducted by undergraduate student Elianna Bruce. L.D. acknowledges financial support from the Israel Scholarship Education Foundation (ISEF). L.A acknowledge financial support by the Israeli Science Foundation (ISF) on the individual research grant number 2107/21, and the New-Faculty Equipment grant number 2108/21; Israeli Ministry of Innovation, Science and Technology 0357-24; BGU’s joint collaboration grant, Engineering Sciences and the Blaustein Institutes for Desert Research, 2025; Pearlston Center at BGU; Israeli Young Academy

## Author contributions

Y.L. and L.A. conceptualized the research. Y.L. and L.A. designed the experiments. Y.S., A.S., and S.N. designed CAD models for 3D printing of inserts and auxiliary parts. Y.L. performed the experiments. Y.L. and L.D. wrote custom codes and carried out all required image analysis. All authors contributed to data interpretation. Y.L. and L.A. wrote the manuscript. L.A. oversaw the project.

## Supplementary material

Supp. movie 1: A narrow insert-based gap closing.

Supp. movie 2: A wide insert-based gap closing.

Supp. movie 3: A Narrow scratch closing.

Supp. movie 4: A wide scratch closing.

Supp. CAD 1: An insert with straight 100 µm wide walls.

Supp. CAD 2: An insert with wavy walls

Supp. CAD 3: A scratcher with 100 µm wide tip

Supp. CAD 4: Shaped islets.

**Supplemental Figure 1.**
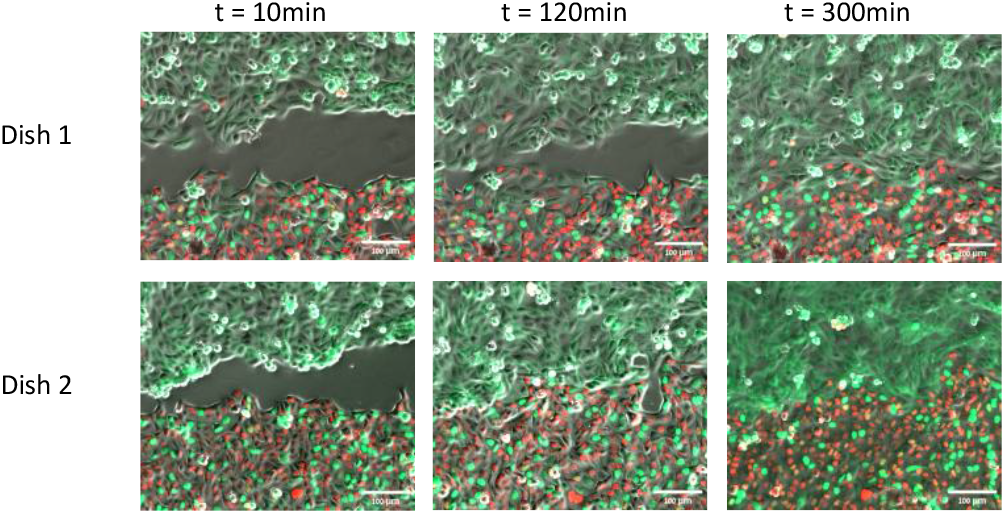
Co-cultures with no intermixing were established with thin-wall inserts that maintained prescribed seeding densities even though cells traversed the narrow gap. The upper region of each panel is seeded with MDCK II cells stably expressing EGFP-tagged ZO-1 (green), while the lower region is seeded with MDCK II cells transfected with the FUCCI system (red/green). Cells were seeded in spatially separated regions using a thin-wall insert, which was subsequently removed to generate a narrow gap between the two populations. Time-lapse imaging reveals the progression of collective migration and interactions at the interface as the gap closes, with no mixing observed before convergence. Importantly, for co-culture purposes, cell area did not increase in short time scales as cells traversed the narrow gap (Supp Figure 2), thereby maintaining the originally prescribed seeding densities. Scale bars: 100 µm.

**Supplemental Figure 2.**
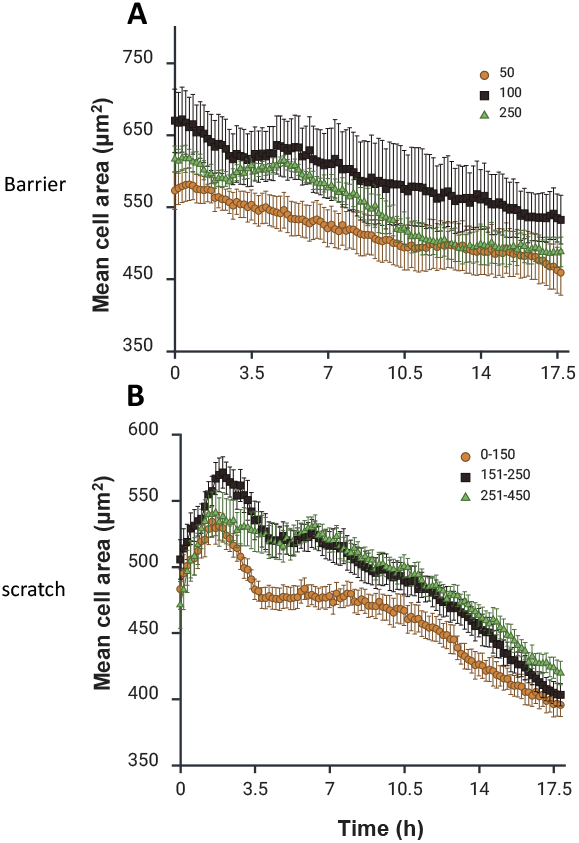
Insert-based assays show a consistent decrease in cell area, indicating continued proliferation throughout, whereas scratch-based assays show an initial increase in cell area, marking expansion rather than proliferation. (**A**) Time course of mean cell area for epithelial sheets closing 50 µm (•), 100 µm (◼), and 250 µm (▲) gaps (mean ± SEM, n = 4, 7, 5, respectively) after insert removal. (**B**) Time course of mean cell area for epithelial sheets closing 0–150 µm (•), 151–250 µm (◼), and 251–450 µm (▲) gaps (mean ± SEM; n = 7, 5, 4, respectively).

## Notes

### Competing Interest Statement

The authors have declared no competing interest.

